# Retrieval Practice Facilitates Judgments of Learning Through Multiple Mechanisms

**DOI:** 10.1101/550731

**Authors:** Xi Chen, Mengting Zhang, Xiaonan L. Liu

## Abstract

Prior studies have shown that predictions of subsequent performance (i.e., Judgments of Learning, JoLs) following practice tests are more accurate than those following re-study. The majority of studies have suggested that retrieval practice allows people to base their predictions on the current retrieval outcomes so that they assign a higher likelihood of remembering the answers with high confidence. We speculated that other information made available through retrieval practice might also be important for JoLs. In the present study, we asked participants to study word pairs and undergo either a practice test or re-study. Two testing formats (cued-recall and multiple choice) were administrated for practice tests in two separate experiments. After each practice trial, participants rated their confidence in the current retrieval accuracy (test) or confidence in acquisition (re-study), followed by a JoL rating where participants predicted their performance in the final test one day later. The results of both experiments showed that the correlation between JoL ratings and the final accuracy was higher for trials practiced with testing. Moreover, using mediation analyses, we found that this high correspondence was only partially mediated by participants’ confidence in practice tests. More importantly, the reaction time of retrieval also significantly mediated the correspondence between JoLs and the final accuracy, suggesting that participants were able to correctly base their JoLs on multiple sources of information that is made available through retrieval practice. We conclude that practice testing benefits JoLs through multiple mechanisms.

## 1. Introduction

Retrieval practice is more beneficial than repeated study: it not only enhances students’ memory performance in a future test (Roediger & Karpicke, 2006) but also helps them make more accurate predictions about future performances (Ariel & Dunlosky, 2011; King, Zechmeister, & Shaughnessy, 1980). The prospective prediction (i.e., Judgments of Learning, JoLs) reflects the quality of metacognitive monitoring of learning and memory, which is critical for efficient learning (Benjamin, Bjork, & Schwartz, 1998; Nelson & Dunlosky, 1991; Nelson, Narens, & Bower, 1990). In an early study (King et al., 1980), participants studied word pairs and underwent either practice test or re-study. Participants then made a JoL rating. Researchers found that participants made more accurate JoLs for the word pairs practiced with a test than for the ones re-studied, suggesting proper monitoring associated with retrieval practice.

Some studies have explored why retrieval improves the accuracy of JoLs (Finn & Metcalfe, 2007; Hertzog, Hines, & Touron, 2013; Serra & Ariel, 2014; Tullis, Finley, & Benjamin, 2013). One account in particular highlights the role of retrieval outcomes: participants assigned a high likelihood of remembering to the items they believed they had been able to answer, and a low likelihood to the ones they could not (Finn & Metcalfe, 2007; 2008). Indeed, in King’s study (King et al., 1980), JoL ratings for successfully recalled items were higher than those not recalled during practice tests. What is largely unknown, however, is whether this subjective belief in retrieval accuracy is the only factor through which retrieval facilitates JoLs.

Dougherty and colleagues (2005) attempted to examine whether JoLs following practice tests simply reflects confidence in the practice retrieval outcome. In their experiment, students were instructed to make two judgments after each practice test: a retrospective judgment about how confident they were that their answer was correct, and a JoL about how confident they would be able to recall the target word in a later test. Results showed that although both judgments were highly correlated, JoLs seemed to be based on separate information from the retrospective judgments because their response distributions were different. This pattern of different response distributions has also been replicated in a recent study (Dougherty et al., 2018), suggesting that additional information may also contribute to JoLs.

Despite the progress in revealing the possible other mechanisms underlying JoLs after practice tests, the majority of research has been focusing on isolating contributing factors, rather than studying JoL as a complex process with multiple bases. Thus, there is still a lack of direct evidence showing what factors may simultaneously contribute to JoLs. Moreover, it leaves the question open as to what specific information made available through retrieval practice promotes the accuracy of JoLs comparing with re-study.

In the present study, we examines whether the high correspondence between JoLs and the final accuracy is fully attributed to participants’ confidence in their retrieval practice performance. We speculate that, in addition to confidence in the retrieval outcome, how easy the answer comes to mind might also serve as a clue for making JoLs. In fact, researchers using general knowledge questions have shown that participants associated the answers that came to mind easily with a higher JoL, even when they were actually harder to remember (Benjamin et al., 1998), suggesting that retrieval ease may be a potent source for JoLs. This study quantitatively examines factors that contribute to more accurate JoLs following retrieval practice, adopting a mediational approach. Specifically, we investigated whether participants’ confidence in retrieval accuracy fully mediated the relationship between the JoLs and the final test accuracy. Moreover, we examined the hypothesis that the reaction time of successful retrievals also served as a clue in making JoLs. We used two different testing formats, cued-recall and multiple-choice, to examine the generalization of results.

## 2. Material and methods

### 2.1. Participants

College students (aged 17-27 years old) participated in this experiment for course credits. Forty-one participants (mean age = 20.5 years, 32 females) completed the experiment with cued-recall. Six of them were excluded due to low accuracy (less than 10%) on the final test. Forty-nine (mean age = 19.7 years, 31 females) completed with the multiple-choice format, ten of which were excluded due to low accuracy. This study was approved by the Ethical Committee, Xiamen University. All participants gave written informed consent in accordance with the Declaration of Helsinki.

### 2.2. Materials and procedures

In this experiment, participants studied Chinese word pairs. A total of 120 word pairs were presented in six lists (for details of stimuli, see Zhang, Chen, & Liu, 2018). During the initial study of each word pair, participants were instructed to memorize word pairs so that they could recall the target when given the cue. After studying a list, each word pair was either tested or re-studied based on the random assignment of conditions in an intermixed order.

We conducted two experiments using two testing formats as the practice test: cued-recall and multiple-choice (Figure 1). During a cued-recall practice test, the cue word was first shown in the center of the screen for three seconds and participants were encouraged to recall the target word. On the next screen, participants were presented with the cue word on the left and a question mark on the right, and were allowed to enter the answer within seven seconds. During a multiple-choice practice test, the cue word was first shown in the center, followed by the presentation of both the cue and four alternatives, which included the correct target word and three synonyms of the target word. Participants had four seconds to enter their answer. No feedback was provided for either condition. A re-study trial presented the intact word pair, and participants would enter the target word within the same time limit.

**Figure 1.**
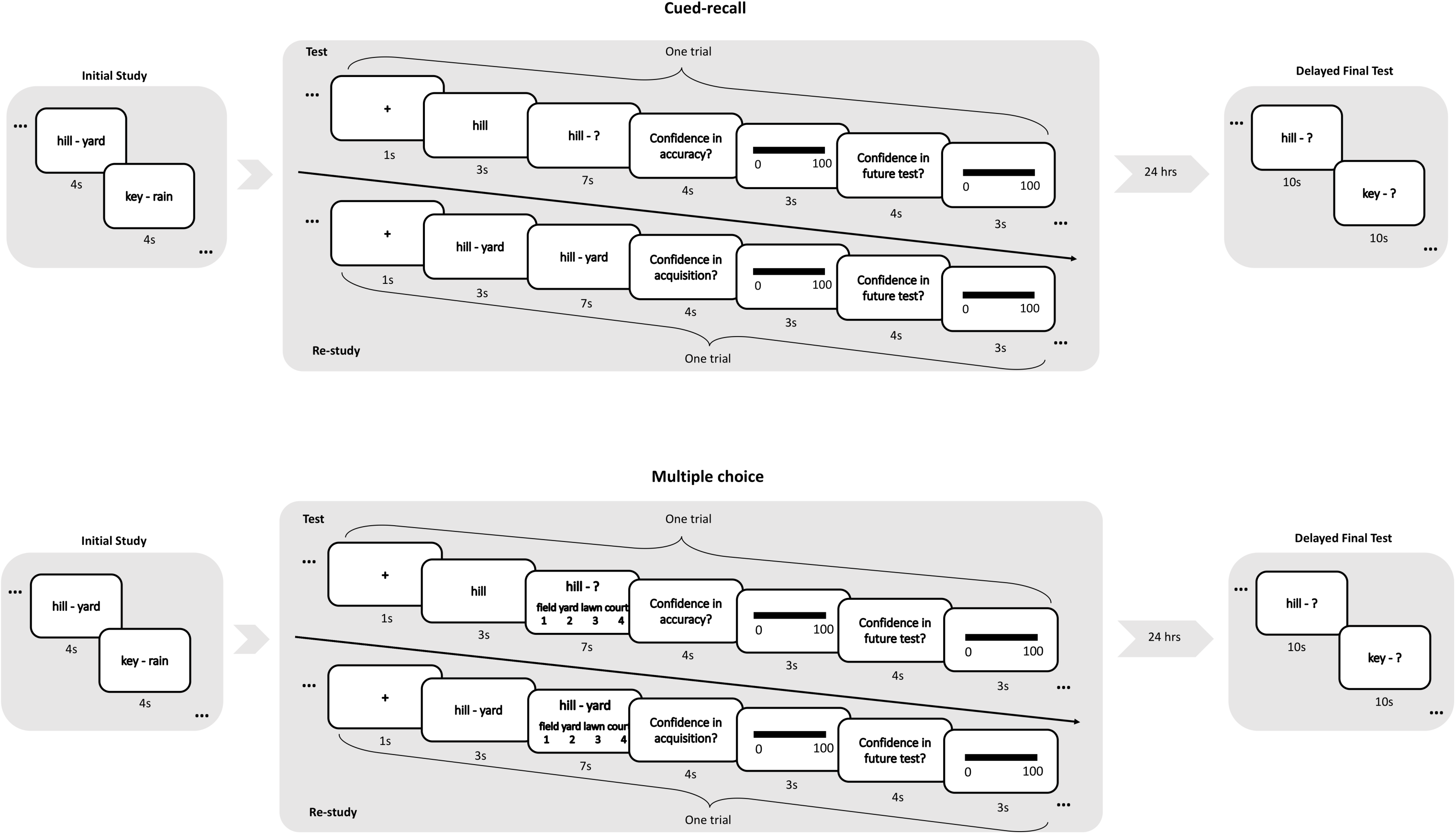
Experimental procedure, using cued-recall and multiple-choice as testing formats in the *test* condition, separately.

After each test or re-study trial, participants rated their confidence in the accuracy of their answer (test trial) or acquisition of the word-pair (re-study trial), on a scale from 0 to 100. Then, participants completed a JoL rating on their confidence, from 0 to 100, that they would be able to remember this word pair for the final test 24 hours later. The final test was administered one day later using a cued-recall test.

## 3. Results

For the cued-recall experiment, participants benefited from practice retrievals and had a higher final accuracy in the *test* condition (*mean*=.332) than in the *re-study* condition (*mean*=.295), *t*=2.08, *p*=.045, as expected. Participants had an average gamma correlation of *r*=.655 between JoLs after *test* trials and final test accuracy, which was significantly higher than the correlation for *re-study* trials, *r*=.338, *t*=8.773, *p*<.001, suggesting that retrieval practice enhanced JoL accuracy. A similar pattern was also found for the multiple-choice experiment: higher final accuracy (*mean*=.282) and JOL correlation (*r*=.488) for *test* than *re-study, mean*=.223, *r*=.380, *t*=3.034, *p*=.004. In addition, the confidence rating on retrieval outcome had high consistency with actual retrieval performance (*r*=.914, *p*<.001 in cued-recall; *r*=.744, *p*<.001 in multiple-choice).

We primarily examined whether participants exclusively relied on the confidence in the retrieval accuracy when making JoLs. Mediation analyses were performed based on mixed model logistic regression using lme4 and RMediation packages in the R Statistical Environment (Bates et al., 2014; MacKinnon et al., 2007; R Core Team, 2016; Tofighi and MacKinnon, 2011). We first modeled the relationship between JoL ratings and the final test performance while including the subject-specific intercepts as the random effect, and then examined the mediation effect of retrieval confidence in this relationship. The analysis revealed that, for the cued-recall experiment, participants’ confidence in the test accuracy significantly mediated JoL’s correspondence with the final performance (Figure 2A), confirming that participants made predictions about the future performance based on their confidence in the practice test. In addition, the mediation effect in the *test* condition was numerically larger than the mediation of acquisition in the *re-study* condition: for *test* trials, indirect/total effect was 79.6%, whereas the ratio is 35.9% for *re-study* trials (Figure 2B), suggesting that subjective confidence in retrieval performance, which is only available after practice tests, was particularly helpful for participants to make better JoLs.

**Figure 2.**
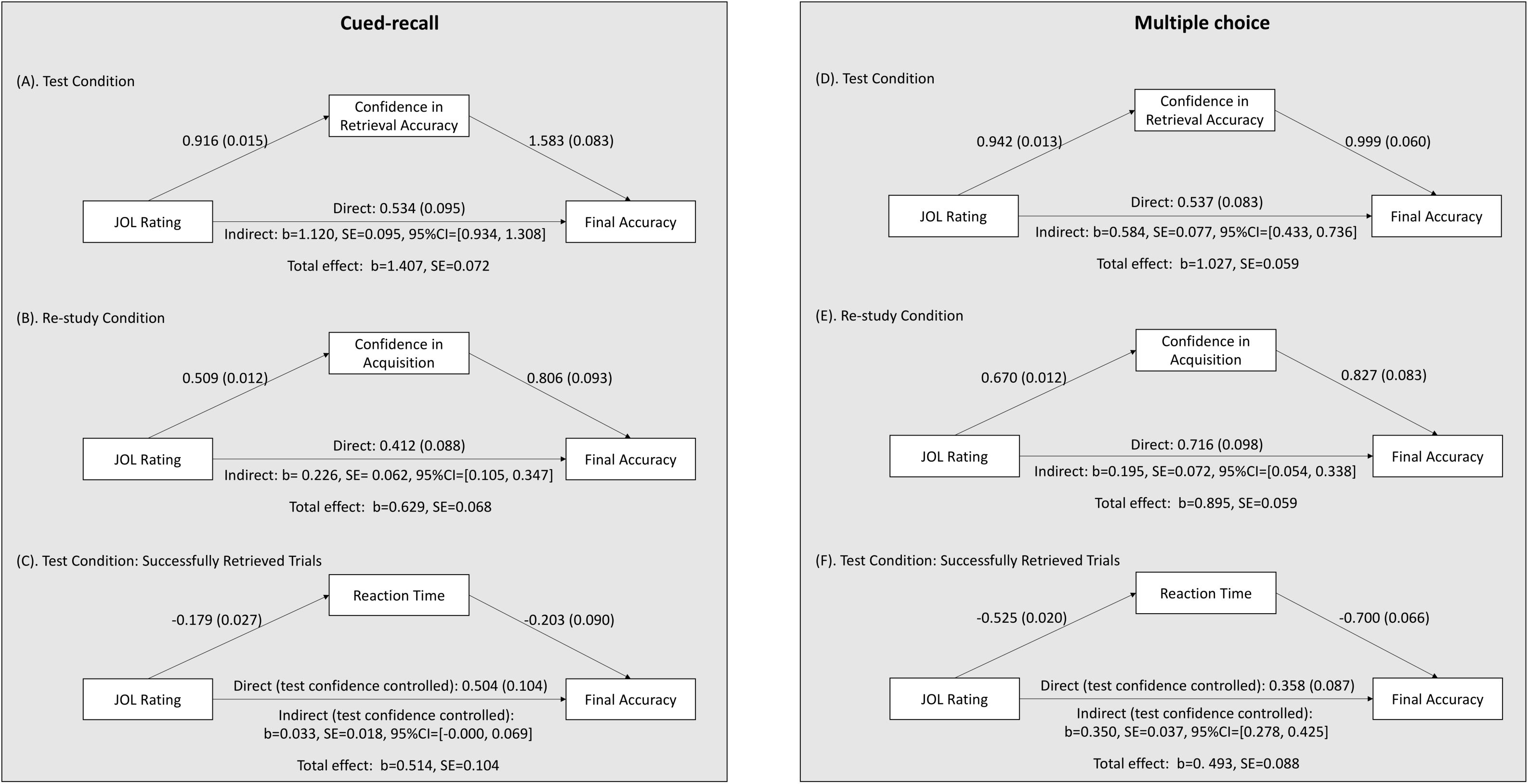
Results from mediational analyses reveal multiple mechanisms underlying the correspondence between JOL rating and final accuracy. In the cued-recall experiment, (A), for the *test* conditions, the mediation of confidence in retrieval accuracy was significant, but did not fully attenuate the relationship between JOL rating and final accuracy. (B) Confidence in acquisition in the *re-study* condition also mediated the relationship between JOL rating and final accuracy for re-studied trials, but the effect is numerically smaller compared to the test condition. (C) After the influence of retrieval confidence taken into account, reaction time of successfully retrieved trials in the test condition also significant mediated the relationship. (D)-(F) The experiment with multiple choice testing format shows consistent results.

Importantly, including the mediator of confidence did not fully attenuate the accuracy of JoL, indicated by a significant direct relationship between JoL ratings and the final test accuracy (Figure 2A), suggesting that other factors may also account for a large portion of the variance in this relationship. We then examined the mediation effect of reaction time for successfully retrieved trials in the *test* condition while controlling for the effect of retrieval confidence. We found that reaction time also significantly mediated the accuracy of JoL beyond the influence of retrieval confidence (Figure 2C), suggesting that participants also relied on how easy the retrieval was when making JoLs, independent of the confidence in retrieval outcomes. We note that this effect was small in magnitude (indirect/total effect = 6.4%), probably because reaction time of a cued-recall response may be affected by the length of the target word.

Finally, we observed the same patterns of results in the multiple-choice experiment (Figure 2D-F), suggesting the generalization of findings for two different practice testing formats. Specifically, the mediation of retrieval confidence was significant (Figure 2D), and the comparison between *test* and *re-study* conditions also revealed a greater mediation effect for the *test* condition (*test*: indirect/total effect = 56.9%, *re-study*: indirect/total effect= 21.8%; Figure 2E). In addition, the effect of confidence in retrieval also did not fully mediate JoL accuracy for multiple-choice trials: reaction time was also a significant mediator after controlling for confidence in retrieval accuracy (Figure 2F; indirect/total effect = 71.0%), confirming that participants used how easy the answer came to mind as a clue for making JoLs and that this strategy helped them make more accurate JoLs.

## 4. Discussion

The goal of the present study was to examine the benefits of practice testing for JoLs and which factors may contribute to more accurate JoLs following retrieval practice. Consistent with previous findings (Finn & Metcalfe, 2007, 2008), we showed that participants rely on the confidence in their retrieval outcomes when making JoLs. Using a mediational approach, we provided direct evidence that this heuristic indeed helps people to give more accurate JoLs. Moreover, we showed that participants’ confidence in their retrieval performance could not fully explain how they were able to achieve high correspondence between JoLs and their final performance, suggesting that other factors may provide additional information that also helps in making better predictions.

A novel contribution of the study is that it complements the understanding of JoLs by showing strong evidence of multiple mechanisms underlying JOLs. Recent studies have discussed this possibility of multiple cues contributing to the processes in making JoLs (Dougherty et al., 2005; Hertzog et al., 2013; Serra & Ariel, 2014). We demonstrated that people also base their judgment on how easy the retrieval is and that this strategy helps them to make better JoLs. This pattern suggests that participants are actively monitoring the processing resources available during retrieval and utilize this information when evaluating and predicting the learning progress.

In fact, researchers have proposed a two-process account (Liu & Reder, 2016; Liu, Tan, & Reder, 2018) for the testing effect, emphasizing a post-retrieval re-encoding process, in addition to the retrieval attempt, that may be related to retrieval monitoring and self-evaluation (Bai, Bridger, Zimmer, & Mecklinger, 2015; Johansson & Mecklinger, 2003). In addition, a prior study using an fMRI showed that retrieval practice, compared with re-study, involved more monitoring and working memory-related brain activity (Liu, Liang, Li, & Reder, 2014). Altogether, the current results that participants rely on retrieval outcomes to make JoLs might reflect the evaluation of the quality of the retrieval attempt process. Moreover, the contribution of retrieval ease might reflect participants’ monitoring of working memory resource used during retrieval and of available resources that could be used for the post-retrieval re-encoding process.

Finally, these findings have important educational implications. Our findings suggest that students may benefit from practice tests prior to an exam, which can not only improve their exam performance, but also allow for better metacognitive monitoring based on their subjective experience during the practice test.

## Author Contributions Statement

XL developed the study concept and design. All authors contributed to data acquisition, analysis, and interpretation of data. XC and XL drafted the manuscript. All authors provided critical revisions of the manuscript for important intellectual context and approved the final version of the manuscript.

